# Obstructed tear duct causes epiphora and precocious eyelid opening due to disruption of Prickle 1-mediated Wnt/PCP signaling

**DOI:** 10.1101/2020.04.17.046383

**Authors:** Dianlei Guo, Jiali Ru, Jiaying Fan, Rong Ju, Kangxin Jin, Hong Ouyang, Lai Wei, Yizhi Liu, Chunqiao Liu

## Abstract

The tear drainage apparatus evolved in terrestrial animals serving as conduits for tear flow. Obstruction of tear drainage causes a range of ocular surface disorders. Hitherto, genetics of tear duct development and obstruction has been scarcely explored. Here we report that a severe *Prickle 1* hypomorph mouse line exhibited epiphora. This phenotype was due to blockage of the tear drainage by the incompletely formed nasolacrimal duct (NLD) and lacrimal canaliculi (CL). Further analysis revealed that the precocious eyelid opening, previously observed in the same type of *Prickle 1* mutants, is also caused by tear duct dysplasia. A comparison of wild type, the *Prickle 1* hypomorph and null mutants revealed a dose-dependent requirement of *Prickle 1* for tear duct outgrowth. As a key component of a set of six Wnt/PCP core proteins, Prickle 1 usually works together with other PCP components. An investigation of expression of Wnt/PCP core genes demonstrated three of the six PCP components in tear duct, supporting the notion of context-dependent organization of PCP protein complexes. Furthermore, expression of *Fgfr2/Fgf10* and *p63* genes, mutations of which are associated with NLD and CL hypoplasia in human, were not altered in *Prickle 1* mutant mice. Lastly, we showed that *Prickle 1* expression in developing tear drainage system is conserved between mouse and human despite anatomical differences. Altogether, the study uncovered how obstruction of the tear drainage could lead to a complex ocular surface disorder, which may have genetic implications in human ocular health.

## Introduction

A major function of the nasolacrimal apparatus in terrestrial animals is to keep the ocular surface from drying. It comprises two systems - the orbital glands and excretory/drainage conduits, each with multiple components. Compromised glandular secretions or tear drainage would lead to a range of ocular surface disorders including the most common dry eye syndrome, underlying genetics of which are barely understood.

The lacrimal canaliculi (CL) and nasolacrimal duct (NLD) are tear drainage conduits opening at orbital and nasal epithelium, and retained in many tetrapods (Frame and Burkat, 2009). The ocular fluid passes across the cornea and conjunctiva and drains into the nasal cavity through CL and NLD (Burling et al., 1991; Thiessen D, 1992). The NLD also possesses absorptive function for tear fluid substances, which may provide feedback signal in tear production, and thus associates with the dry eye disorder (Paulsen et al., 2002a; Paulsen et al., 2002b). Phylogenic variations of drainage ducts exist extensively among species (Rehorek et al., 2015; Rehorek et al., 2011; Rossie and Smith, 2007; Tamarin and Boyde, 1977). Anatomically, human and rabbit NLD share many similarities (Frame and Burkat, 2009; Paulsen et al., 2002b) (de la Cuadra-Blanco et al., 2006) despite their different origins, with the former originating from ectodermal lacrimal groove and the latter from subcutaneous region of the lower eyelid (Rehorek et al., 2011). Thus, rabbit is considered a suitable model for human tear duct study from the anatomical perspective. Mouse NLD is suggested to develop similarly as humans based on a scanning electron microscopy study (Tamarin and Boyde, 1977). However, little is known about the anatomy, genetics and development of the mouse NLD.

Clinically, congenital nasolacrimal duct obstruction (CNLDO) was estimated to be present in up to 20% of newborn infants (Vagge et al., 2018; Wallace et al., 2006), often causing epiphora with dacryocystitis and conjunctivitis. Genetic risk factors for CNLDO are poorly known. There are several scattered reports of CNLDO in syndromic diseases, suggesting existence of the genetic elements for CNLDO (Foster et al., 2014; Inan et al., 2006; Jadico et al., 2006a; Jadico et al., 2006b; Kozma et al., 1990; Rohmann et al., 2006; van Genderen et al., 2000). For instance, mutations in FGF signaling components (FGF10, FGFR2 and FGFR3) lead to Lacrimo-auriculo- dento-digital (LADD) syndrome exhibiting hypoplastic NLD and puncta, often in conjunction with conjunctivitis (Rohmann et al., 2006). In accordance with these findings, ankyloblepharon-ectodermal dysplasia-cleft lip/palate (AEC) syndrome caused by mutations in p63, a transcription factor for *Fgfr2* (Ferone et al., 2012), has lacrimal duct obstruction overlapping with LADD, in which tear outflow obstruction can occur with agenesis of NLD and CL (Allen, 2014). Nonetheless, genetic etiology of non-syndromic CNLDO has been uncovered, and currently CNLDO genetics is barely known.

The Wnt/PCP pathway has been shown to be involved in ocular development and diseases recently. Particularly, a mutation in *Prickle 1* in mice causes ocular surface inflammation and precocious eyelid opening (Guo et al., 2019), the primary cause of which has yet to be identified. Additionally, a delayed embryonic eyelid closure was also observed in *Prickle 1* mutant mice (Guo et al., 2018). Prickle is one of a set of six proteins (Frizzled, Disheveled, Vangl, Prickle, Diego, and Flamingo) that executes PCP signaling (Jenny and Mlodzik, 2006), which is critical for a variety of tissue morphogenesis (Wallingford et al., 2002; Wallingford and Harland, 2002). As such, depending on genetic background and allelic variations, *Prickle 1* mutants display pleiotropic defects from gastrulation (Tao et al., 2009) to organogenesis (Liu et al., 2014; Yang et al., 2013). We recently noticed the *Prickle 1* heterozygous compound mutants having watery eyes (epiphora). We suspected that tear duct might be obstructed in the mutants to block tear drainage. In the current report, we confirmed this speculation and present additional evidence that *Prickle 1* was required for normal tear duct development, malformation of which is the trigger for epiphora, precocious eyelid opening and ensuing ocular surface pathogenesis. Further investigation demonstrated that *Prickle* gene family is also expressed in human tear drainage system during development, suggesting a possibility that *Prickle 1* might be an important genetic contributor to CNLDO.

## Materials and methods

### Mice and genotyping

Animal husbandry and experimentation were conducted in strict adherence to the Standards in Animal Research: Reporting of In Vivo Experiments (ARRIVE) guidelines, with approval from Animal Care and Use Committee (ACUC), Zhongshan Ophthalmic Center, Sun Yat-sen University. The severe *Prickle 1* hypomorph mutant mouse line used in this study was generated by crossbreeding a *Prickle 1* gene-trap mutant allele (*Prickle 1^a/+^*) to a straight knockout allele (*Prickle 1^b/+^*) (Guo et al., 2019; Liu et al., 2014; Liu et al., 2013). Mouse genotyping was conducted as described previously (Liu et al., 2014; Liu et al., 2013). Mouse strains used are mixed genetic backgrounds from C57BL/6 and Sv129.

### Human embryos

The human embryonic materials were provided by the Guangzhou Women and Children’s Medical Center. The specimens were obtained from miscarriages and verified no congenital malformations. All human study protocols were reviewed and approved by the Institutional Review Board of Guangzhou Women and Children’s Medical Center.

### Histology

For fresh frozen sections of mouse heads, the postnatal mice (P1, P5, P8) were sacrificed by decapitation and adult mice were sacrificed by cervical dislocation. The dissected heads were directly embedded in OCT (SAKURA, Cat. 4583, USA) and frozen in liquid nitrogen and store at -80 0C freezer. Sections were cut at 15 μm for all immunostaining purposes except for 3- Dimensional reconstruction, in which 30 μm sections were cut.

For paraffin eyeball sections, the eyeballs including the intact eyelids were dissected and fixed in 4% paraformaldehyde for 24h at 4℃ (Guo and Liu, 2019). The samples were washed 3 times in PBS (phosphate buffered saline), dehydrated through a series of alcohols and 3 times of xylene, then embedded in paraffin and sectioned using microtome (Leica RM 223, Wetzlar, Hesse-Darmstadt, Germany). Hematoxylin (H) (Cat. G1002, Servicebio, China) and Eosin (E) (Cat. G1004, Servicebio, China) staining followed manufacturer instruction.

For prefixed frozen sections, mouse heads were bisected and PFA-fixed as stated before. After PBS washes, tissues were submerged in 30% sucrose overnight for cryoprotection, embedded in OCT and store at -80 0C before sectioning.

### Immunohistochemistry

For frozen sections and fresh frozen sections, immunostaining followed a standard protocol. Briefly, tissue sections were blocked with 10% donkey serum with 0.1% triton in PBS (PBST) for 30min at RT, then incubated with primary antibodies at 4℃ for overnight. After washed with PBST, sections were incubated with the fluorescent dye-conjugated second antibodies for 1hr at room temperature, washed with PBST and mounted with Fluoromount-G (Southern Biotech, Birmingham, AL, USA). Antibodies used in this study are anti-E-cad (ab11512, Abcam) and anti-p63 (ab124762, Abcam).

### In situ hybridization (ISH)

Digoxigenin (11277073910-Roche) (DIG)-labeled sense and antisense RNA riboprobes were prepared by in vitro transcription from T7 (5’ - TAATACGACTCACTATAGGG-3’) and T3 (5’-AATTAACCCTCACTAAAGGG-3’) promoter-tagged PCR fragment from each gene using corresponding T7 and T3 RNA polymerase (T3:M0378S; T7:M0251S, BioLabs). Primers used for PCR are listed in Supplemental table 1. ISH was performed using a protocol that was described by Jensen and Wallace (Jensen and Wallace, 1997).

### Tear fluid collection, protein gel and western blot analysis

Tears were induced by intraperitoneally injection of pilocarpine (300 μg/kg weight) and collected from the eyelid margin into Eppendorf tubes using a 0.5 μl micropipette five minutes after injection. Same volume of tears from wild type and the mutant mice were used for running a 10% SDS-PAGE. Protein profiles were visualized by Coomassie brilliant blue staining for 2h at RT followed by destaining with 40% methanol and 10% glacial acetic acid. For western blot, protein samples were blotted onto PVDF membranes by a wet transblot system (Mini-Protein Tetra, Biorad) using standard protocol recommended by the manufacturer. After blocking with 5% fat-free milk, membranes were incubated with primary antibody against mouse lactoferrin (07-685, Millipore) and then peroxidase-conjugated secondary antibody IgG (M21002S, Abmart, China). Chemiluminescent images were taken by FluorChem R (Proteinsimple).

### Imaging

Fluorescence microscopy images were obtained using Zeiss confocal microscope (Zeiss LSM880, Zeiss, Oberkochen, Germany) and Imager.Z2 equipped with ApoTome (Zeiss, Oberkochen, Germany). H&E and ISH images were acquired by Imager.Z2.

### 3D reconstruction of tear duct

Images taken from microscopes were aligned manually by Photoshop software according to anatomical features of each section. NLD structure was traced on each section and imported to NIH ImageJ software for 3D- processing (with 3D viewer plugins) with properly set image and pixel depth and adjusted image coordinates.

### Quantification and statistics

For measuring the missing length of NLD, coronal sections from P1 wild type, severe *Prickle 1* hypomorph and null mutant heads were subjected to H&E staining. The first section displaying NLD in both wild type and the mutant mice was designated as “zero NLD length”, respectively. Missing NLD in the mutants was calculated as vertical distances between section at “zero NLD length” and the section exhibiting most similar anatomical structures to that of wild type at “zero NLD length” (Missing NLD length =section thickness × No. of sections). 6 animals were used for each genotype. Student t-test was used to detect power of significance.

## Results

### Tear duct dysplasia in a severe *Prickle 1* hypomorphic mutant led to epiphora

A severe hypomorphic *Prickle 1* mutant that is compound heterozygous for the null and hypomorphic alleles was generated (see Materials and Methods, Supplemental Fig. 1A) (Guo et al., 2019; Guo et al., 2018). We estimated that Prickle 1 was expressed less than 25% of WT levels from western blots using limb and brain tissues (Supplemental Fig. 1B, C). This severe hypomorph exhibited epiphora at adulthood (Fig. 1A, B). We suspected that the tear drainage was obstructed, prompting us to examine the tear duct histology. Transverse fresh frozen sections from the nasal proximity of the mouse head (Fig. 1C) were subjected to H&E staining (Fig. 1D-O). Sections from wild type and the mutant mice of the same anatomical structures were compared (Fig. 1D-F & G-I compared with J-L & M-O, respectively). Absence of the distal NLD was observed in all examined mutant animals (n=3) (Fig. 1J-O). We next examined CL, which normally branches out from the proximal NLD and opens at the inner eyelids. Oblique sections parallel to the facial plane were prepared (Fig. 1P-W). By E-cadherin staining, openings of both upper and lower canaliculus to the front canthus were detected in wild type mice (Fig. 1P, Q), but not in the mutants (Fig. 1T), even though all other corresponding structures could be found in both wild type and the mutant mice (Fig. 1R, S compared to U and V, respectively). Thus, the missing ends of the CL blocking tear drainage explains the epiphora phenotype.

**Figure 1.**
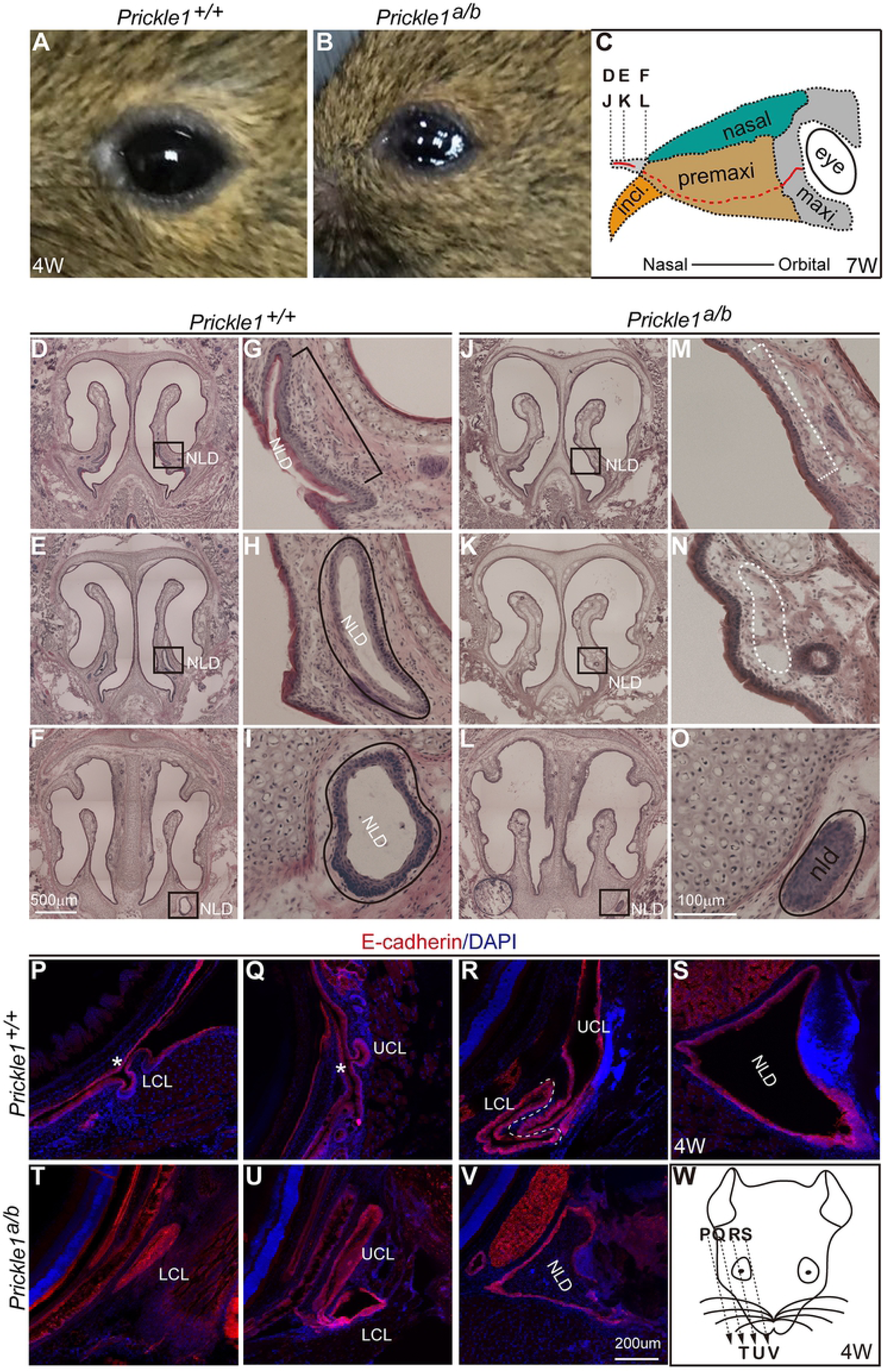
Tear duct histology and epiphora in severe *Prickle 1* hypomorph mutants. Abbreviations are applied to all figures unless otherwise noted. Fresh frozen sections were subjected to H&E and immunohistochemistry. (A) Normal cornea surface. (B) Mutant cornea surface exhibiting epiphora. (C) Section positions in (D-F) and (J-K). (D-F) Wild type nasal sections. Boxed areas are nasolacrimal duct (NLD). (G-I) Magnified from (D-F). Bracket and lines indicate NLD. (J-L) Mutant nasal sections positionally correspond to that of wild type in (D-F). Boxed areas are magnified in (M-O). (M-O) Dashed bracket and lines indicate presumptive NLD. (P-V) E-cadherin staining revealed NLD and canaliculi. (P-S) Representative sections of wild type tear duct from different positions. Sectioning direction is from temporal to nasal as illustrated in (W). UCL, upper canaliculus; LCL, lower canaliculus; Asterisk, canaliculi openings at eyelids. (T-V) Mutant tear duct sections from temporal to nasal. (W) Illustration of cutting direction and sectioning positions.

### Tear duct dysplasia coincided with pooling of tear fluid under the eyelid in *Prickle 1* compound mutants

The missing nasal NLD and orbital CL suggested a developmental abnormality of the drainage system. We thus examined early postnatal ages of mice to inspect the consistency of this phenotype. An examination of NLD using H&E histology demonstrated similar missing nasal end of the mutant NLD at P8 (Fig.2 A-C), while retaining the medial part (Fig. 2D-E). Immunostaining of E-cadherin revealed missing orbital proximal openings of the canaliculi (Fig. 2F, G right panels compare with the left), whereas the remaining parts of CL connecting with NLD were generally preserved (Fig. 2H, I right panels compare with the left). These data are consistent with the findings presented in adult mutant mice (Fig. 1). Surprisingly, we observed remarkable accumulation of ocular fluid under eyelid, apparently pressuring the lid to become thinner and nearly open (Fig. 2K). This observation thus explains the mutant precocious eyelid opening previously found by Guo (Guo et al., 2019), and led us to further investigate mutant ocular fluid production in time series.

**Figure 2.**
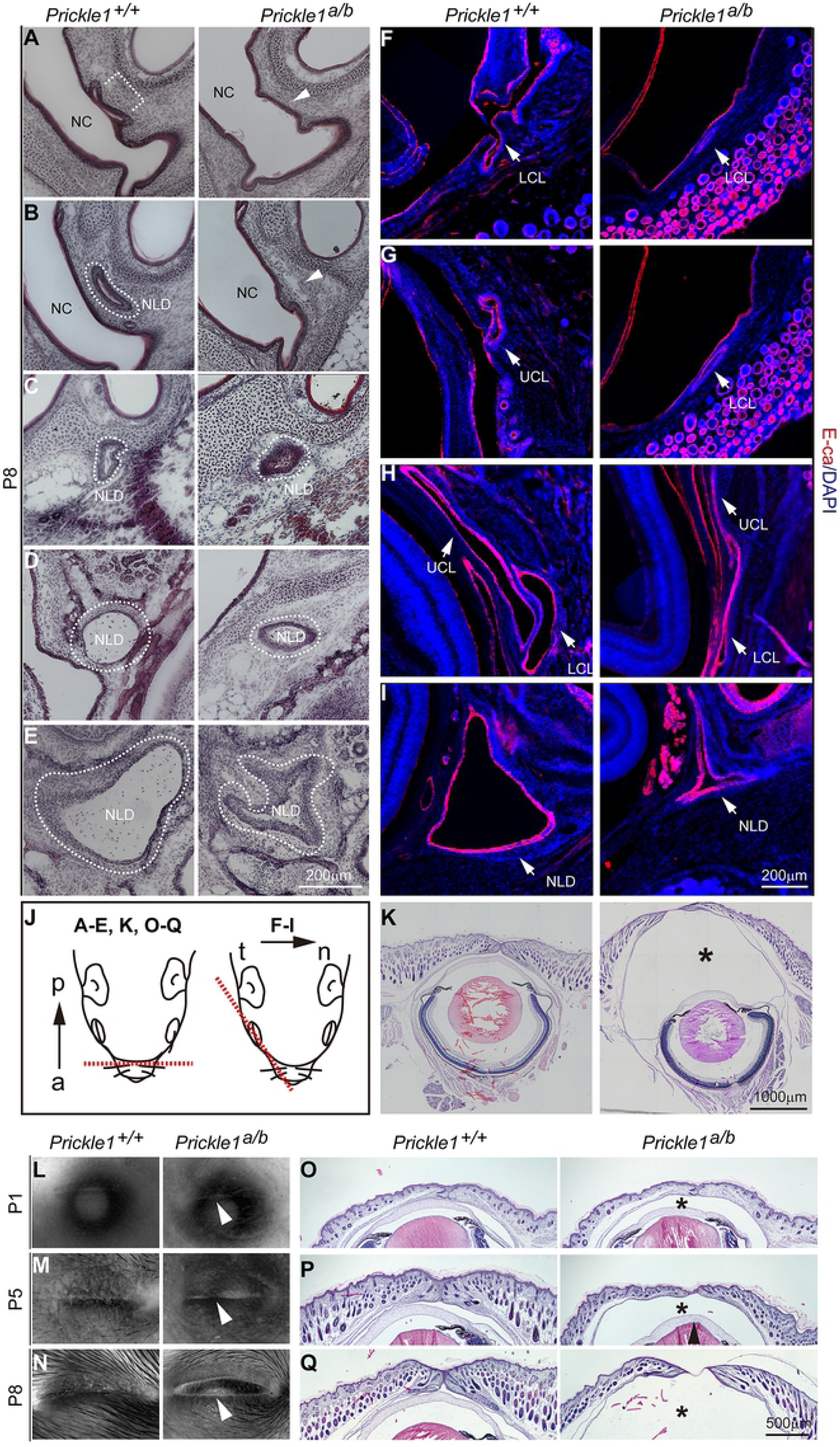
Tear duct obstruction, tear fluid retention and ruptured eyelid in *Prickle 1* hypomorphic mutants. (A-E) H&E-stained fresh frozen sections. NC, nasal cavity; Arrows and lines indicated NLD positions. (F-I) E- cadherin staining revealed NLD and CL. Note the openings of lacrimal canaliculi in lower eyelid (F) and upper eyelid (G) of the wild type P8 mice were missing in the mutants. The furthest extended mutant UCL on section is shown in right panel of (H). Arrows point to different parts of the tear duct. (J) Section direction for (A-E) (left) and (F-I) (right). a, anterior; p, posterior; t, temporal; n, nasal. (K) Whole-eyeball coronal sections at P8. The asterisk indicates mutant ocular fluid accumulation under the eyelid. (L-M) Eyeballs viewed from top. Arrows point to the mutant eyelid. (O-Q) H&- stained coronal sections of eyelid.

Top view of P1 eyeballs did not show differences between the mutant and wild type eyelid (Fig. 2L). Bulging and smoother surface of the mutant lid was detected at P5 (Fig. 2M, right panel), and stretched eyelid was obvious at P8, when the lid junction nearly broke (Fig. 2N, right panel). On H&E stained sections, mutant increased ocular fluid was reflected by larger space between the cornea and eyelid, even obvious at P1 (Fig. 2O), which progressively expanded until eyelid was ruptured (Fig. 2P, Q). These data suggested that obstruction of tear drainage caused by the malformed tear duct resulted in the mutant ocular fluid retention and eyelid rupture.

### Mutant ocular fluid is tears and ocular glands are largely normal

Should the accumulated mutant ocular fluid be due to obstructed tear duct, it must be tears in nature. To confirm this speculation, we compared the mutant ocular fluid with the wild type tears at P8 before eyelid open. Pilocarpine was used to stimulate tear production for P8 wild type mice, because insufficient amount of tear can be collected due to their draining down to naris. On the contrary, there is no need to stimulate the mutants because of continuously accumulated fluid tank under the eyelid. Tears or mutant ocular fluid from P8 mice were drawn with syringes, and same volumes were run on SDS denature gel. Protein profiles were comparable between the mutant and wild type mice at P8, as predicted (Fig. 3A). We next stimulate tears production of both wild type and the mutant mice at 6-week old and performed the same analysis as we did for P8. Similar results were observed--no qualitative differences were detected on SDS-PAGE gel (Fig. 3A). Consistently, lactoferrin/transferrin, a major component of tears, was also expressed similarly in wild type tears and the mutant ocular fluid (Fig. 3B). We therefore concluded that the mutant ocular fluid was primarily tears, and the mutant lacrimal gland function was largely normal.

**Figure 3.**
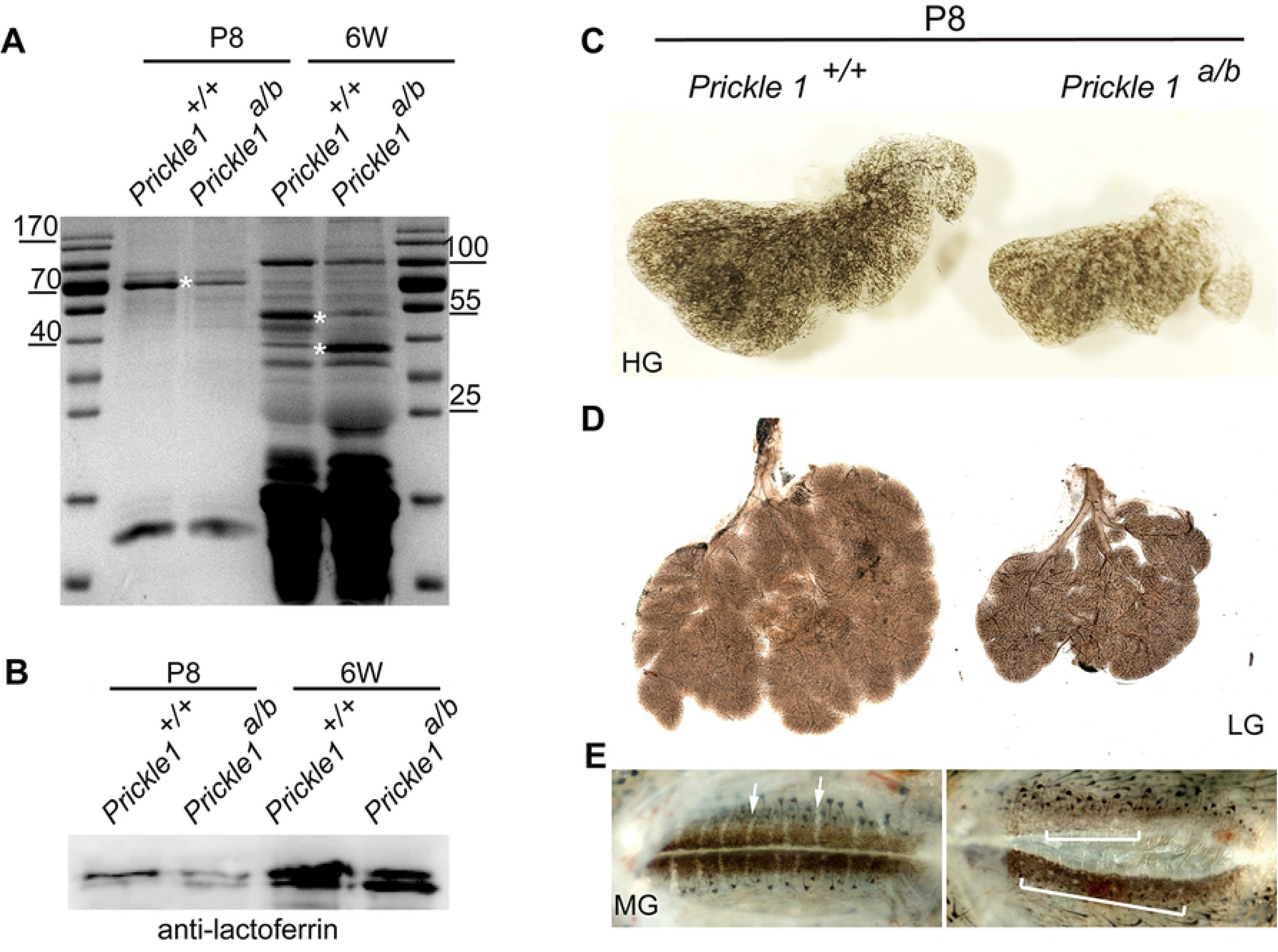
Accumulated ocular fluid in *Prickle 1* hypomorphic mutants is primarily tears. (A) SDS-page profiling the mutant and wild type tears at P8 and 6-week old (6W). Asterisks indicate differential intensities of similar molecular shifts in the mutant and wild type tears. (B) Immunoblotting of the mutant and wild type tears using anti-transferin antibody. (C) Wild type and the mutant P8 Harderin glands (HG). (D) Lacrimal glands (LG). (E) Meibomian glands (MG) indicated by arrows and brackets respective for wild type and the mutant mice.

The accumulated tear fluid in the mutants might also result from excessive production of the ocular glands in combination with the dysplastic tear duct. Notwithstanding, gross examinations of the ocular glands at P8 showed general normal morphology of the mutant Harderin and lacrimal glands with the same number of lobes (Fig. 3C, D). The gland sizes were smaller matching with the body sizes. Additionally, Meibomian gland was aberrant in the mutants (Fig. 3E), which might be due to compression by the accumulated tears. These results indicated otherwise normal if not compromised gland secretions.

### Dose-dependent requirement of *Prickle 1* on tear duct formation

In general, *Prickle 1* null alleles have more severe phenotypes than hypomorphic or null/hypomorph compound alleles. This could be partially reflected from the fact that hypomorph/null *Prickle 1* compound mutants (*Prickle ^a/b^*) survive normally, whereas null mutant (*Prickle ^b/b^*) mice dies within ~24 hours after birth (Liu et al., 2014). We thus reasoned that *Prickle 1* null alleles might also give rise to a more severe tear duct phenotype. We compared tear duct of different genotypes at P1, when *Prickle ^b/b^* mice were still alive. On H&E-stained sections, wild type NLD completely reached nasal cavity and opened at P1 (Fig. 4A). Consecutive sections demonstrated regionally irregular tube shape of NLD with apparent stratified epithelium (Fig. 4B-E). In contrast, *Prickle 1 ^a/b^* NLD was about 300 μm (section vertical thickness) away from its nasal destination (Fig. 4F-J). The distal most NLD was missing in the *Prickle 1^b/b^* mutants, which is about 1.5 mm (Fig. 4K-O). Quantification of the vertical NLD length by counting sections demonstrated significant dose-dependent missing of NLD on disruption of *Prickle 1* gene (Fig. 4P-Q).

**Figure 4.**
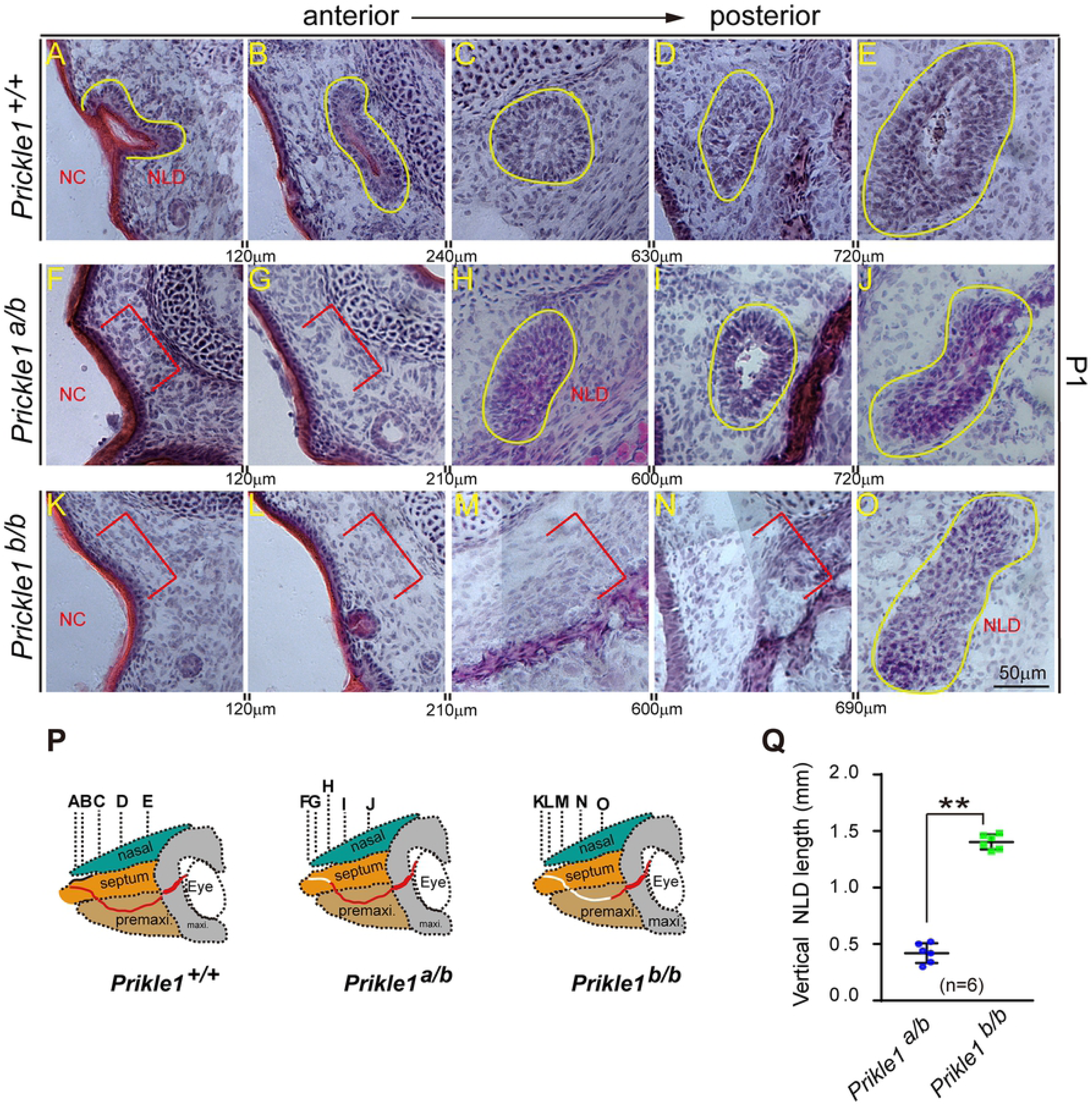
*Prickle 1* dose-dependent loss of nasolacrimal duct. (A-O), H&E-stained fresh-frozen sections. Yellow lines indicate existing NLD, red brackets indicate missing of NLD. Interval distances are shown between each panel. (A-E) Wild type NLD at P1. (F-J) *Prickle 1* hypomorphic mutant duct with null (*Prickle 1^b^*) and hypomorph (*Prickle 1^a^*) alleles. (K-O) *Prickle 1^b/b^* null mutant duct. (P) Schematic illustration of dose-dependent loss of NLD (white lines) in *Prickle 1* hypomorphic compound heterozygous mutants. (Q) Quantification of missing NLD of *Prickle 1^a/b^* and *Prickle 1^b/b^* mutants. Missing NLD was quantified as vertical distances (number of sections × section thickness) from presumptive nasal ends to sections that first appeared NLD. Student *t-test* was performed to detect significance.

### Expression of Fgf and Wnt/PCP signaling components in mouse tear duct development

Because Fgf and Wnt/PCP pathways are known to be extensively involved in duct morphogenesis, we investigated their expression in early age of tear duct development. We focused on embryonic day E11, when growing tear duct could be identified from the eyelid and conjunctiva expressing epithelial marker, p63 (Fig. 5A-P). *Fgf10* was expressed surrounding tear duct (Fig. 5A, B) complementary to *Fgfr2*, which was expressed in tear duct proper (Fig. 5C, D). In contrast, *Fgfr3* was not expressed in tear duct despite its strong expression in lateral ventricles (Supplemental Fig. 2A-D). A systemic investigation of expression of Wnt/PCP components found that among 19 Wnts, *Wnt 6* was highly expressed in tear duct and skin epithelium (Fig. 5E, F, Supplemental Fig. 2M), whereas other *Wnts* were only weakly expressed or did not have distinct expression pattern (Table 1, Supplemental Fig. 2). Out of 10 Frizzled receptors, *Fz3* and *Fz6* were weakly expressed in initial parts of the tear duct (Fig. 5G-J, Table 1, Supplemental Fig. 3). Two of the three PCP atypical cadherins, *Celsr 1* and *Celsr 2* were expressed in initial and full tear duct, respectively (Fig. 5K-N, Table 1, Supplemental Fig. 4A). Interestingly, none of *Dvl*, *Vangl* families or *Inversin* were expressed in developing tear duct with a distinct pattern (Supplemental Fig. 4). As expected, *Prickle 1*, but not other family members, are restricted to tear duct indicated by an antisense probe against *eYFP* reporter gene (Fig. 5O, P, Supplemental Fig. 5). The reporter expression was verified by *Prickle 1* antisense probe (Fig. 5Q, R (E11.5)). Taken together, the expression of Fgf signaling components was consistent with the NLD and CL phenotypes observed in human patients bearing *Fgf10/Fgfr2* mutations, and the six PCP core genes did not seem to work together all the time.

**Figure 5.**
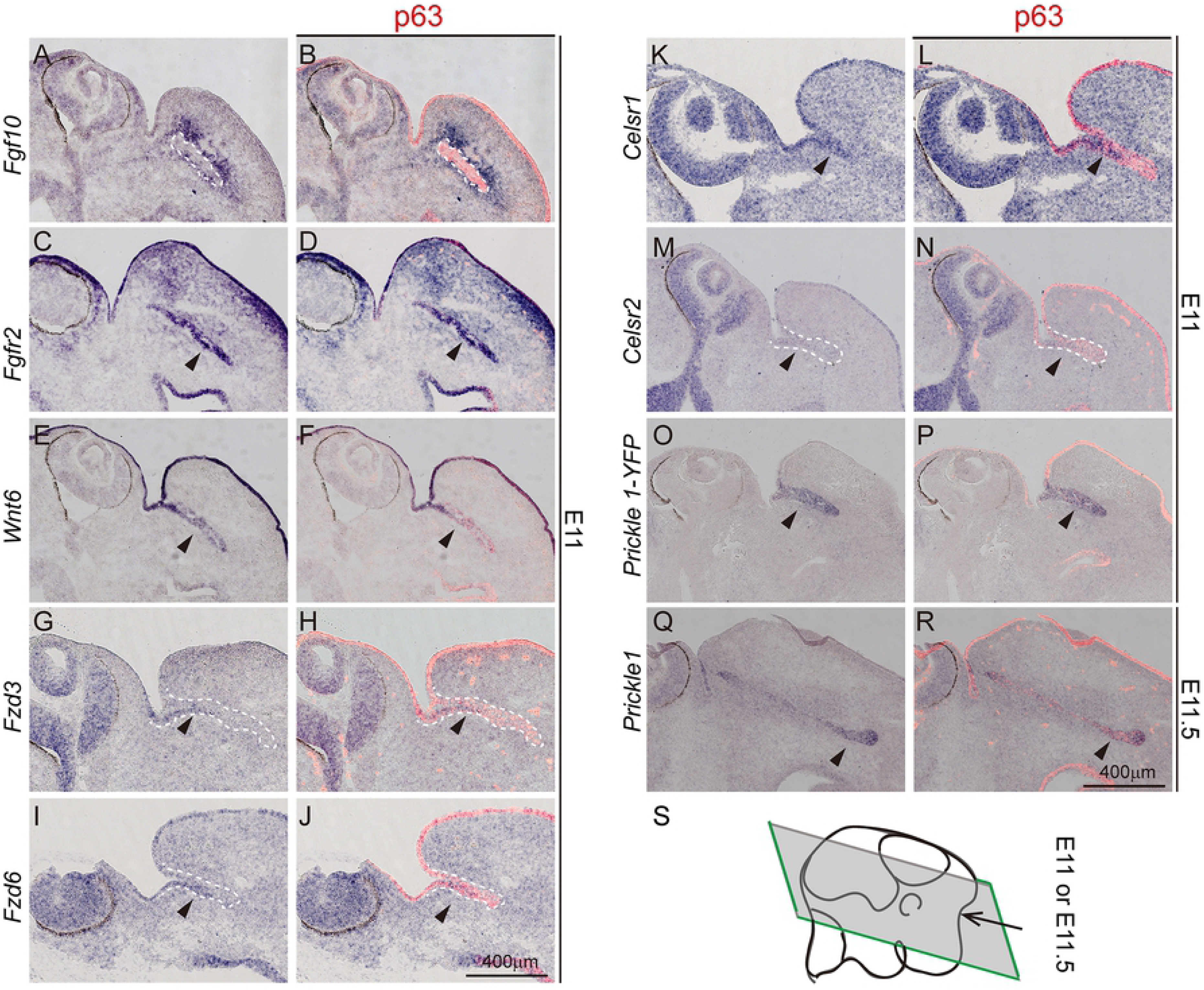
Expression of *Wnt/PCP* and *Fgf* signaling components in developing tear duct. All panels are in situ hybridization (purple) followed by p63 immunostaining (red). Dashed lines demarcate tear duct, arrows indicate gene expression. (A, B), *Fgf10*. (C, D), *Fgfr2.* (E, F), *Wnt6*. (G, H), *Fzd3*. (I, J), *Fzd6*. (K, L), *Celsr1*. (M, N), *Celsr2*. (O, P), *Prickle 1*. (Q), Schematic illustration of a horizontal sectioning plane through developing tear duct at E11.

**Table 1:** A summary of expression of Wnt/PCP components in developing mouse tear duct at E11.

| Wnt/PCP family | Tear duct | Surrounding tear duct | Uniformly or no expression |
| --- | --- | --- | --- |
| *Wnt* | *Wnt1, 4, 6* | *Wnt11, 5a* | *Wnt2, 2b, 3, 3a, 5b, 7a, 7b, 8a, 8b, 9a, 9b, 10a, 10b, 16* |
| *Prickle* | *Prickle 1* | - | *Prickle 2, 3, 4* |
| *Frizzled* | *Fzd 3, 6* | *Fzd 10* | *Fzd 1, 2, 4, 5, 7, 8, 9* |
| *Dvl* | - | - | *Dvl 1, 2, 3* |
| *Celsr* | *Celsr1, 2* | - | *Celsr3* |
| *Vangl* | - | - | *Vangl1, 2* |
| *Inversin* | - | - | *Inversin* |

### Disruption of *Prickle 1* did not alter p63, *Fgf10*, *Fgfr2* or *Wnt6* expression

The distinct expression of *Wnt 6*, p63, *Fgf10*, and *Fgfr2* in the tear duct prompt us to ask whether their expression would be altered in *Prickle 1* mutant mouse. To address this, we used wild type and *Prickle 1* null mutant embryos for examination. The outgrowth of the tear duct was apparently stunted at the examined age of E11 (Fig. 6A-H). In situ hybridization showed no remarkable difference in expression of p63 (Fig. 6A, B), *Fgf10* (Fig. 6C, D) and *Fgfr2* (Fig. 6E, F) between wild type and the *Prickle 1* mutant mice. *Wnt 6* expression also remained similarly in wild type and the mutant tear duct (Fig. 6G, H). The results thus suggest that genetically, *Prickle 1* is either downstream of, or parallel with *Fgf* signaling during tear duct growth.

**Figure 6.**
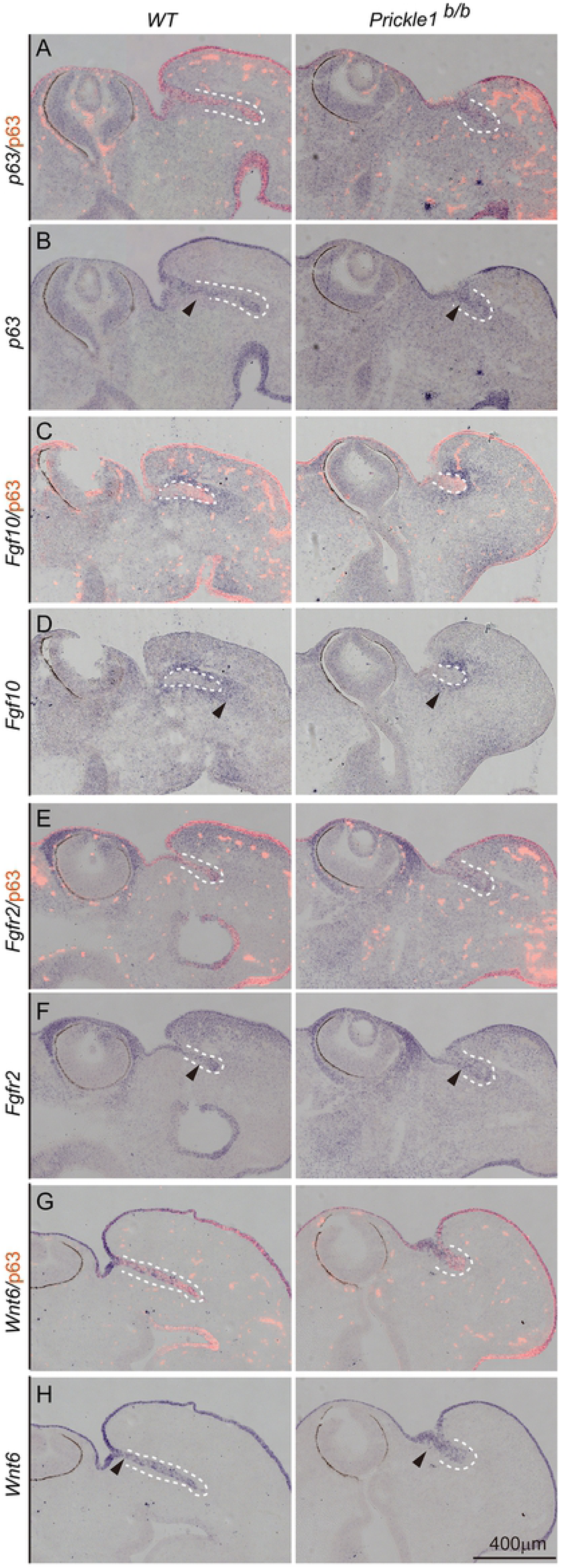
Expression of *Fgfr2, p63, Fgf10 and Wnt 6* in *Prickle 1* mutant tear duct. Dashed lines demarcate tear duct, arrows indicate gene expression. All panels are in situ hybridization (purple) followed by p63 immunostaining (red). (A, B), *Fgf10*/p63. (C, D), *Fgf10*/p63. (E, F), *Fgfr2*. (G, H), *Wnt 6*.

### Expression of *PRICKLE* gene family in human tear duct development

We predicted that the function of *Prickle 1* in tear duct development might be conserved among mammals. And it is especially of interest to see this in humans to explore potentials of *Prickle 1* mutants being a suitable disease model. We therefore investigated expression of *PRICKLE* gene family in developing human tear duct. We collected gestational weeks 8 (GW8) old embryos, when lacrimal groove has formed by the fusion of lateral nasal and maxillary processes (de la Cuadra-Blanco et al., 2006). By in situ hybridization followed by immunostaining of p63, *PRICKLE 1* is highly expressed in developing human tear duct (Fig. 7A-F). Interestingly, unlike in the mouse, human *PRICKLE* 2, 3, and 4 were also expressed in developing human tear duct with a distinct pattern, though weakly (Fig. 7G-L). Furthermore, all of *PRICKLE* genes exhibited unusual dotted expression pattern, prompting us to examine the developmental path of the human tear duct. Using p63 as an epithelial marker, we reconstructed images from all embryonic sections comprising fetal tear duct at GW8. On 3D developing duct, multiple tubular branches were found (Fig. 7M-P, Supplemental video). These branches generally extended in vertical plane since top view of the 3D image missed most of them (Fig. 7O, P). The pattern might reflect unsynchronized pinching off of the regional lacrimal groove epithelial cells, collectively and independently migrated downward.

**Figure 7.**
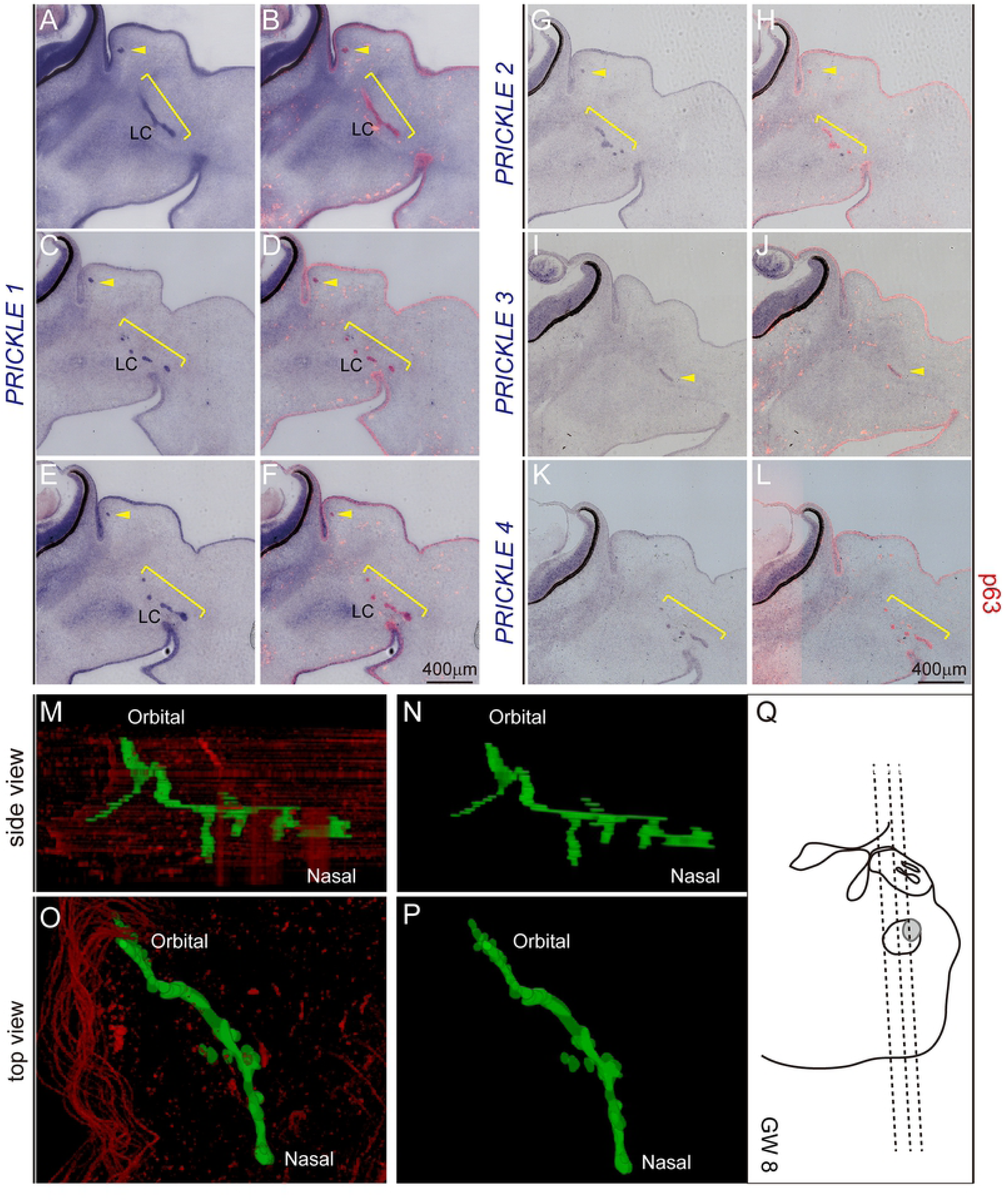
Expression of PRICKLE family in developing human tear duct. (A-L), All panels are in situ hybridization (purple) followed by p63 immunostaining (red) at gestation week 8 (GW8). (A), (C) & (E) are three consecutive sections with 30 μm intervals. (B), (D) & (F) are same sections respective to (A), (C) & (E). (A-F). *PRICKLE1/*p63. (G, H) *PRICKLE2/*p63. (I, J) *PRICKLE3/*p63. (K, L) *PRICKLE4/*p63. (M-P), 3-D reconstruction of embryonic tear duct with p63 staining drawn as green. (M, N), Side/lateral view of tear duct. (O, P) Top view of tear duct. (Q), Schematic illustration of section orientation.

## Discussion

Obstruction of tear drainage would lead to a range of ocular disorders often with epiphora, keratoconjunctivitis and dacryocystitis. Despite its importance, little attention has been paid to this system, probably because: 1) tear duct obstruction is not a life-threatening disease, therefore insufficient genetic data were collected in clinics; 2) it is entirely embedded in complex bony and cavernous tissues, restricting its accessibility to research; and 3) no suitable animal models are available for human studies. In attempt to fill these blanks, the current study demonstrate how obstruction of tear drainage could lead to a chain of ocular surface disorders using a genetically engineered mouse model potentially useful for human-related diseases. We further extend our investigations to previously unexplored genetic determinants of the tear duct, providing a framework for future understanding developmental biology and diseases of the drainage system. Our work additionally provides a potential tear drainage-related human disease model, and offering opportunities to explore novel biological functions of PCP.

The hypomorph/null *Prickle 1* compound mutant mice exhibited epiphora leading us to suspect whether obstructed tear drainage may occur. Histology study confirmed such speculation. Interestingly and surprisingly, it turns out that previously identified precocious eyelid opening and ocular surface pathogenesis (Guo et al., 2019; Guo et al., 2018) are all due to the malformed tear duct in *Prickle 1* mutant mice. In normal mice, lacrimal gland already starts to secrete tears at P1, which drain through the tear conduits to naris. In *Prickle 1* mutant mice, however, although tears are normally produced as in wild type before eyelid opening, it fails to drain out. The continuous production of tear fluid without drainage builds up pressure against the closed eyelids constraint, eventually forcing them to open. The ruptured eyelid debris then falls back to the corneal and conjunctival surface further eliciting keratoconjunctivitis (Guo et al., 2019). It is remarkable that the complex spectrum of ocular surface phenotypes in the *Prickle 1* mutant mice is all triggered by malformed tear duct.

Although apparently coevolved, lacrimal gland develops much later than tear duct (Garg and Zhang, 2017). *Prickle 1* seems to have a more specific role in tear duct morphogenesis than in ocular glands. In fact, morphology the ocular glands are largely normal in *Prickle 1* mutants, and tear components do not change much either. The primary role of *Prickle 1* in tear duct could also be reflected from the dosage-dependent phenotypic manifestation of the mutant NLD. Anatomically, NLD could be roughly divided into three segments along the head axis: orbital/proximal, bony/middle and narial/distal (W. J. Hillenius et al., 2005; Rehorek et al., 2015). The *Prickle 1* heterozygous compound mutants miss only partial distal NLD, whereas null mutants miss more. The graded phenotypic severities suggest that the *Prickle 1* is highly likely a key determinant for tear duct morphogenesis.

Because *Prickle 1* is a key component of Wnt/PCP core protein complex, we investigated expression of all known Wnt/PCP components. Additional to confirmation of *Prickle 1* expression in tear duct, we found that not all PCP core complex genes are expressed together with *Prickle 1*. This further suggests that tissue context-dependent combinations of PCP core members coordinate morphogenesis of different tissues. Among the 19 *Wnts*, only *Wnt 6* is highly expressed in tear duct. However, whether it transduces signaling through μ-catenin dependent or PCP pathway is yet to be investigated.

Three Fgf signaling components, *Fgfr2, 3*, *Fgf10* and p63, a upstream transcription factor for *Fgfr2*, are known to be involved in human LADD syndrome (Allen, 2014; Ferone et al., 2012; Rohmann et al., 2006) with obstructed NLD and CL. Consistently, *Fgfr2*, *Fgf10* and p63 are all expressed in mouse tear duct or surrounding tissues, except for *Fgfr3*. It is appealing to know whether Wnt/PCP and Fgf signaling would genetically interact. However, we did not observe remarkable changes in expression of the examined Fgf components in *Prickle 1* mutants, implying that Prickle 1- mediated PCP functions either parallel or downstream of Fgf signaling. Notably, *Fgf10/Fgfr2* pair also plays an essential role in lacrimal gland development (Garg and Zhang, 2017), which is barely affected in *Prickle 1* mutant mice. Therefore, it is likely that Fgf and Wnt/PCP operate on different aspects of tear duct development. Nonetheless, whether Wnt/PCP pathway is downstream of Fgf signaling is yet to be clarified.

The similar expression of *PRICKLE1* in both human and mouse tear duct implies its evolutionally assigned function in this organ. Yet unlike in mouse, other *PRICKLE* members were also weakly expressed. Human developing tear duct also exhibits more branches from the epithelial cord, which are not seen in the mouse. The meaning of these differences is biologically interesting to be unearthed. Additionally, human eyelid opens well before birth. Whether there would exists eyelid condition similar to that of *Prickle 1* mutant mouse are difficult to address. Nevertheless, nearly 20% of newborn infants bear CNLDO, often with epiphora, dacryocystitis and conjunctivitis, which are all manifested in *Prickle 1* mutant ocular surface. Combined with the conserved expression of *Prickle 1* in lacrimal apparatus, these observations suggest that common molecular machineries may contribute to mouse and human tear duct diseases. Although *Prickle 1* is crucial to many tissues, the observed dosage effect of *Prickle 1* in mouse together with expression of multiple members of *PRICKLE* family in human may provide redundancy to allow individual survival yet with CNLDO related-ocular diseases. With such speculation, *Prickle 1* mutant mouse could be a useful animal model to be explored for human tear duct diseases.

## Acknowledgements

We thank Drs. Tiansen Li from the National Eye Institute for providing valuable suggestions in preparation of the manuscript. The authors thank Drs. Tiansen Li for critical reading the manuscript and helpful comments. We thank lab members Shujuan Xu, Shanzhen Peng, Shiyong Zhu, for technical supports to the work.

## Supplemental Figure Legends

**Supplemental Figure 1.(A)** Genomic structure and mutant alleles of *Prickle 1* gene (from Liu et al, 2014). *Prickle 1*^a^ (hypomorphic) and *Prickle 1*^**b**^ (null) alleles were bred together to produce *Prickle 1*^a**/b**^ compound mutant mice, designated as severe *Prickle 1* hypomorphic mutants in this study. (**B**) Immunoblot of tissue extracts from E13.5 embryonic hindpaws was probed with a customized polyclonal Prickle 1 antibody. Prickle 1 was not detectable in *Prickle 1*^**b/b**^ and only weakly expressed in *Prickle 1*^a**/b**^ mutants (from Liu et al., 2014). (**C**) Weak Prickle 1 expression in P1 brain cortex of severe *Prickle 1* hypomorphic mutants. Arrows points to expected Prickle 1 band. Based on immunoblot, Prickle 1 expression was estimated as less than 20% of the wild type levels.

**Supplemental Figure 2. Expression of *Fgfr3* and Wnt family members.** Embryos are at E11. Dashed lines indicate tear duct. P63 immunostaing (pink) was performed after in situ hybridization (ISH). (A) Sectioning plane. (B) *Fgfr3* was expressed in lateral ventricle, but not in tear duct. (C) p63 immunostaining on *Fgfr3* ISH. Boxed areas are magnified in D. (E-W) Expression of all 19 *Wnt* family members. Only micrographs of tear duct region are shown.

**Supplemental Figure 3. Expression of *Frizzled* family members.** (A-J) Horizontal sections. Schematic section plane is illustrated in Figure 5S. Dashed lines indicate tear duct in each panel. Same experiments were performed on all panels as described in Supplemental Figure 2.

**Supplemental Figure 4. Expression of *Celsr 3*, *Dvl* and *Vangl* family members and *inversin*.** Same experiments were performed on all panels as described in Supplemental Figure 2. (A) *Celsr 3*/p63. (B-D) *Dvl* (1-3)/p63. (E, F) *Vangl* (1, 2)/p63. (G) *Inversin*/p63.

**Supplemental Figure 5. Expression of *Prickle* family members.** Same experiments were performed on all panels as done for Supplemental Figure 2. (A) *Prickle 2*/p63. (B) *Prickle 3*/p63. (C) *Prickle 4*/p63.

**Supplemental video caption:** 3D-reconstruction of human developing embryonic tear duct based on ISH images. Green indicates the tear duct.

**Supplemental Table 1:** In situ hybridization primers used in this manuscript.

